# Immunometabolic rewiring in long COVID patients with chronic headache

**DOI:** 10.1101/2023.03.06.531302

**Authors:** Suan-Sin Foo, Weiqiang Chen, Kyle L. Jung, Tamiris Azamor, Un Yung Choi, Pengfei Zhang, Suzy AA Comhair, Serpil C. Erzurum, Lara Jehi, Jae U. Jung

## Abstract

Almost 20% of patients with COVID-19 experience long-term effects, known as post-COVID condition or long COVID. Among many lingering neurologic symptoms, chronic headache is the most common. Despite this health concern, the etiology of long COVID headache is still not well characterized. Here, we present a longitudinal multi-omics analysis of blood leukocyte transcriptomics, plasma proteomics and metabolomics of long COVID patients with chronic headache. Long COVID patients experienced a state of hyper-inflammation prior to chronic headache onset and maintained persistent inflammatory activation throughout the progression of chronic headache. Metabolomic analysis also revealed augmented arginine and lipid metabolisms, skewing towards a nitric oxide-based pro-inflammation. Furthermore, metabolisms of neurotransmitters including serotonin, dopamine, glutamate, and GABA were markedly dysregulated during the progression of long COVID headache. Overall, these findings illustrate the immuno-metabolomics landscape of long COVID patients with chronic headache, which may provide insights to potential therapeutic interventions.

## INTRODUCTION

The unprecedented global pandemic of Coronavirus Disease 2019 (COVID-19) has caused over 665 million infections and 6.7 million deaths (as of 11 Jan 2023)^1,2^. COVID-19 is triggered by the severe acute respiratory syndrome coronavirus 2 (SARS-CoV-2), a new single-stranded RNA betacoronavirus that relies on receptors such as angiotensin-converting enzyme 2 (ACE2) to enter host cells in order to establish infection^3^. Patients who contracted COVID-19 have a wide spectrum of symptoms, from being asymptomatic to developing severe symptoms such as acute respiratory distress syndrome and multiple organ failures^4^. Although most COVID-19 patients recovered from the respiratory disease within a few weeks, there is mounting clinical evidence that these post-COVID patients continue to experience persistent symptoms including headache, fatigue, shortness of breath, loss of smell and taste, depression, anxiety, cardiovascular damage, memory loss as well as joint pain. These patients are often called ‘long-haulers’ and diagnosed with a condition termed post-acute sequelae of SARS-CoV-2 infection (PASC). Among these lingering symptoms, post-COVID headache is one of the most common neurologic symptoms reported in over 70% of active SARS-CoV-2 infection^5,6^, while approximately 40% of COVID-19 patients developed chronic headaches which persisted up to three months post-infection^7–9^. Patients with post-COVID headaches experienced severe debilitating migraine-like pressure pain that spreads throughout the skull and persists for several months, leading to high morbidity^10^.

The association between primary headaches and inflammation has been well reported, yet the pathophysiology of COVID-19-related secondary headaches is still unclear^11^. COVID-19 is considered a multi-organ disease that is attributed by systemic inflammation^12,13^. In fact, the contribution of immunometabolism to the regulation of immune response and inflammation during viral infections, such as human immunodeficiency virus (HIV) and Dabie bandavirus (DBV), has been widely reported^14–16^. Recently, a metabolomics sera analysis of COVID-19-positive patients showed alterations in metabolites involved in arginine and tryptophan metabolisms which correlated tightly with circulating inflammatory cytokines such as IL-6, IP-10 and M-CSF, highlighting the role of perturbed host metabolisms in response to SARS-CoV-2 infection^17,18^.

The neuro-invasion of SARS-CoV-2 has been shown to result in brain vascular inflammation, but the long-term impact of COVID-19 neurological sequalae is still unknown^19,20^. Interestingly, longitudinal effects of COVID-19 showed significant reduction in brain size and cognitive decline, suggesting that neuro-inflammation triggered by SAR-CoV-2 infection interferes with neuronal functions and disrupts neurotransmitter pathways ^21^. Indeed, a dysregulation in the neurotransmitter systems such as serotonin, dopamine and glutamate has been implicated in migraine pathogenesis and pain regulation^22^. Although SARS-CoV-2 is expected to trigger inflammation, immune activation, and dysregulation of metabolites^23,24^, the biological factors associated with persistent and disabling chronic headaches specifically in the context of long COVID remains to be fully characterized.

To address the immuno-metabolomics landscape of post-COVID headaches in ‘long haulers’, we performed a longitudinal multi-omics profiling of peripheral blood specimens of PASC patients over the course of chronic headache progression. (Fig. 1A). Ultimately, identifying key factors that drive post-COVID headaches could provide insights for potential immunotherapies.

**Fig. 1.**
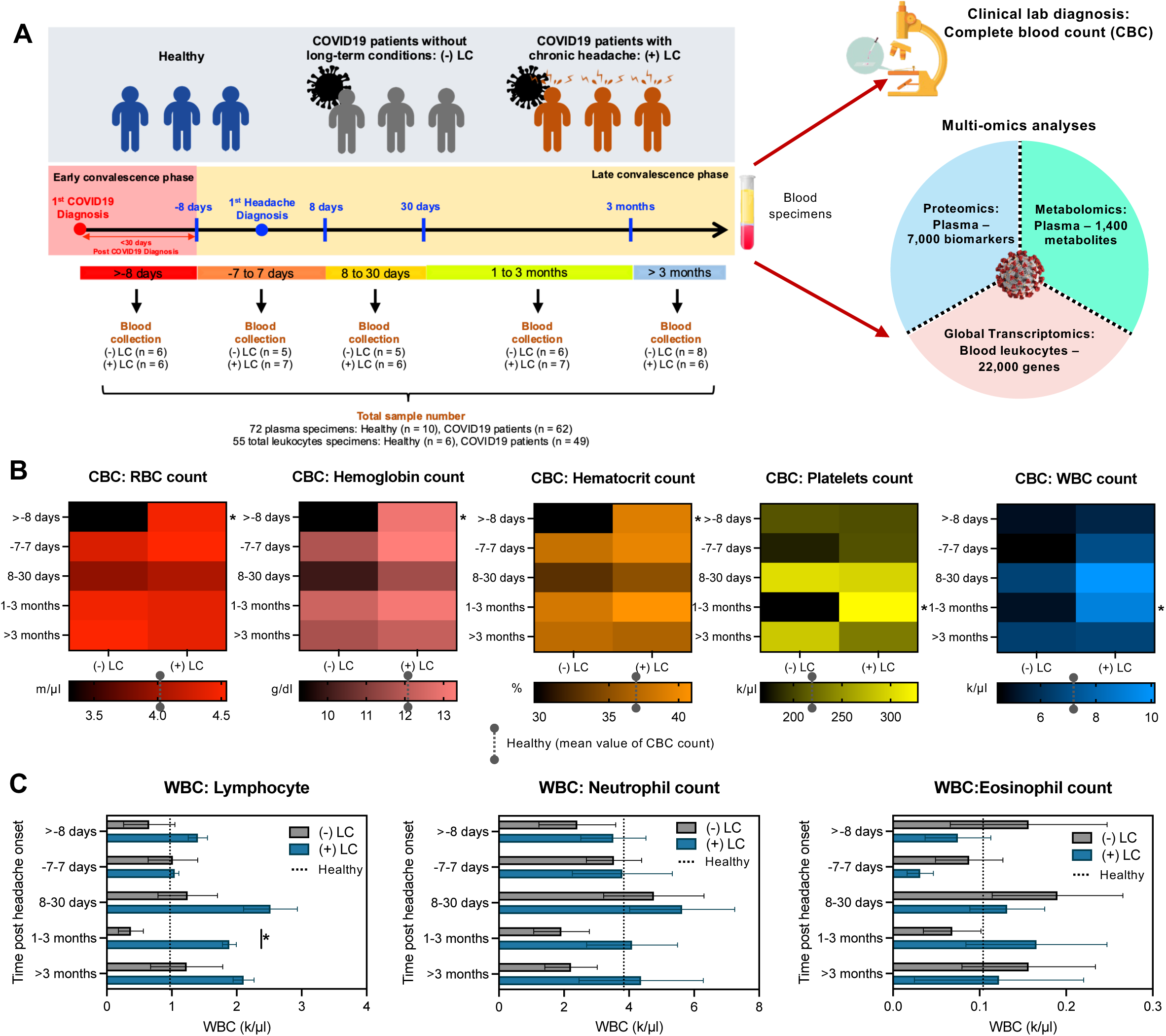
Overview of clinical cohort design and clinical parameters of patients’ blood specimens. (A) Overview of the long COVID – headache study of COVID-19 patients experiencing post-acute sequelae of prolonged headache. 72 blood specimens were collected over 5 time points post SARS-CoV-2 confirmed diagnosis. Clinical lab diagnoses were performed on these blood specimens. Plasma portions were subjected proteomics and metabolomics profiling, and packed cells were subjected to transcriptomics profiling. (B) Complete blood count (CBC) of the various blood components. (C) Cell counts of various white blood cells (WBCs) subsets. For (B) and (C), data are presented as mean ± SEM, using Welch *t*-test. \**P* < 0.05.

## RESULTS

### Overview of clinical and immuno-metabolic features of COVID ‘long haulers’

In this study, we obtained retrospective clinical specimens collected during the COVID19 pandemic period between May 2020 to October 2021, from a clinically well-characterized patient cohort enrolled in the Cleveland Clinic BioRepository (CC-BioR) and the Cleveland Clinic COVID Registry. Blood cells (n = 55) and plasma (n = 72) specimens were collected from age-matched healthy controls (n = 6 to 10), COVID19 patients (n = 49 to 62) with long COVID [(+)LC] associated with chronic headache conditions and without long COVID [(−)LC] (Fig. 1A and Table S1). All COVID19 patients were PCR-confirmed for SARS-CoV-2 infection, blood specimens were collected over the course of post-acute phase of COVID19 disease, including the following time points post headache onset: (i) >-8 days, (ii) −7 to 7 days, (iii) 8 to 30 days, (iv) 1 to 3 months and (v) >3 months. All blood specimens collected were subjected to clinical lab diagnosis for complete blood count (CBC) or multi-omics analyses which included bulk RNAseq transcriptomics, SomaScan Discovery proteomics (~7,000 biomarkers) and Metabolon metabolomics (~1400 metabolites) (Fig. 1A). Interestingly, CBC tests revealed significant reduced levels of red blood cells (RBC) in (−)LC patients compared to (+) LC patients at >-8 days prior to headache onset, which correlated with the reduced levels of hemoglobin and hematocrit (Fig. 1B). In contrast, (+)LC patients exhibited pronounced elevated levels of white blood cells (WBC) and platelets at later phase of COVID19 illness between 1 to 3 months post headache onset compared to (−)LC group (Fig. 1B). Specifically, lymphocyte counts were markedly increased in (+)LC patients at 1 to 3 months post headache onset compared to (−)LC patients, while no significant differences were observed for other subsets of WBC including neutrophil and eosinophil counts (Fig. 1C).

### Overview of multi-omics analyses of long COVID19 patients with chronic headache

To date, the exact mechanism underlying the persistence of long COVID symptoms remains elusive, particularly in the aspect of chronic headaches. Using multi-omics approach, we seek to evaluate the immuno-metabolomics alterations during the different phases of long COVID associated chronic headache symptom (Figs. 1A and 2A). Overall t-SNE plots of the multi-omics analyses revealed distinctively broad alterations of immuno-metabolomic profiles of COVID19 patients, shifting away from healthy controls, while overlapping profiles between (−)LC and (+)LC groups were clearly observed (Figs. 2B, 2E and 2H). To delineate the specific biomarkers associated with (+)LC chronic headache symptom, we employed an analysis strategy as outlined in Fig. 2A. Briefly, immuno-metabolic profiles encompassing transcriptomics, plasma proteomics and plasma metabolomics were first compared (i) between Healthy and (−)LC, as well as (ii) between Healthy and (+)LC. The number of significantly altered (upregulated or downregulated) genes, proteins and metabolites from each time points were identified. These biomarkers of genes, proteins and/or metabolites were further categorized into (−)LC only group, (+)LC only group, or common group, as represented in the Venn diagram (Fig. 2A).

**Fig. 2.**
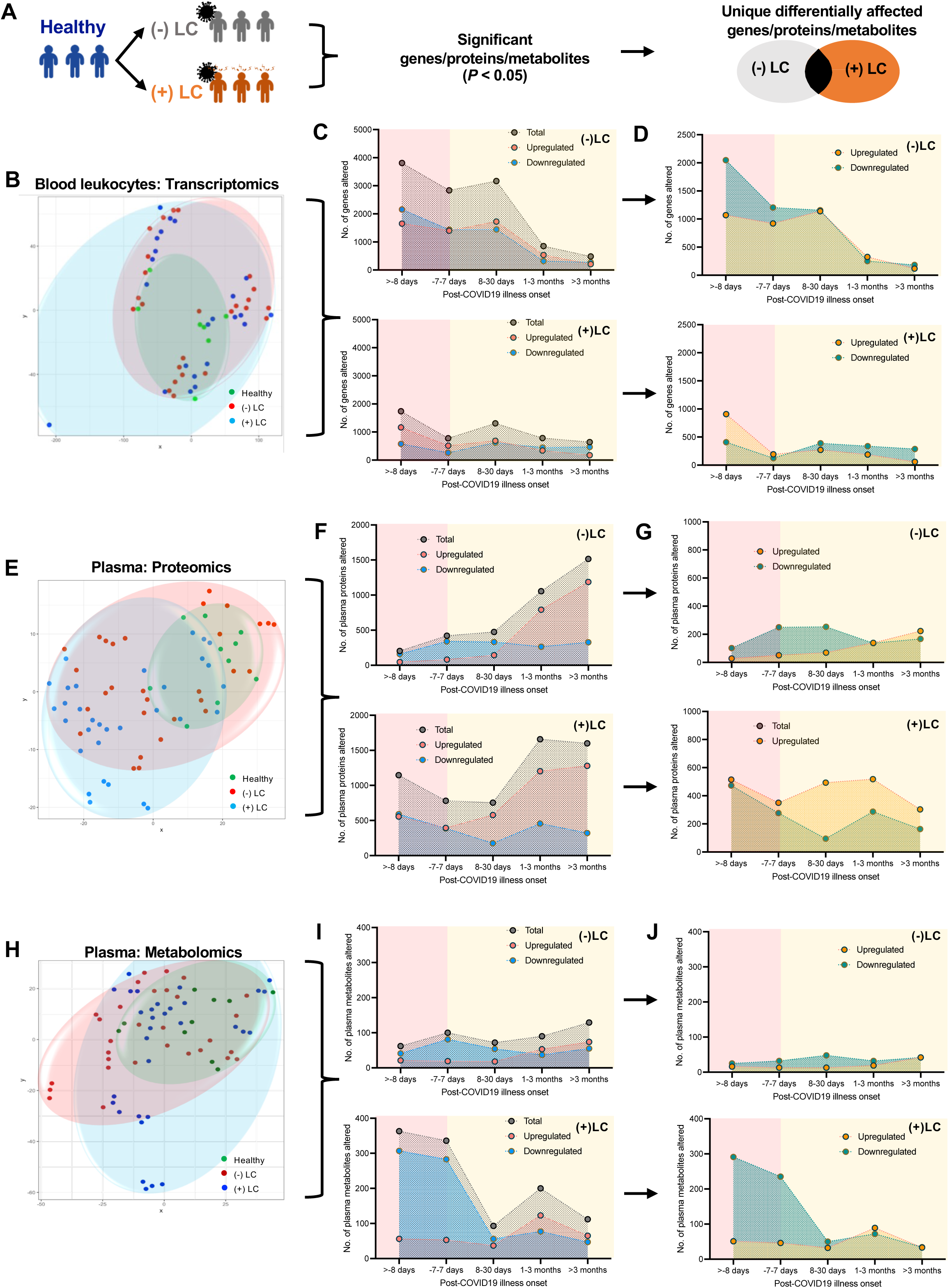
Overview of multi-omics analyses reveal immunometabolic alterations in long COVID patients with chronic headache. (A) Schematic representation of analyses strategy for multi-omics dataset. (B, E, H) t-SNE plots of transcriptomics (B) profiles of blood cells, proteomics (C) and metabolomics (D) profiles of plasma specimens from healthy, (−)LC and (+)LC groups. (C, F, I) No. of genes/proteins/metabolites significantly affected (*P* < 0.05) in (−)LC and (+)LC groups relative to healthy controls. (D, G, J) No. of genes/proteins/metabolites uniquely affected in (−)LC or (+)LC group only, excluding overlapping host factors that are affected in both groups.

RNAseq analysis of the blood leukocytes revealed higher transcriptomics changes in (−)LC patients compared to (+)LC patients, especially at the earlier time points within the first month following an acute COVID illness (Fig. 2C and 2D). Specifically, at >-8 days post-COVID19 illness onset, (−)LC patients exhibited pronounced immune suppression with 2047 genes downregulated and 1069 genes upregulated. In contrast, (+)LC patients exhibited an opposing trend skewing towards immune activation with 907 genes upregulated and 408 genes downregulated (Fig. 2D). In contrast to transcriptomics profiles, plasma proteomics profiles of both (−)LC and (+)LC exhibited considerable alterations of plasma proteins at the later timepoints, post COVID19 illness onset from 1 month onwards (Fig. 2E and 2F). Upon elimination of the overlapping plasma proteins that were significantly altered in both groups of COVID19 patients, (+)LC patients clearly demonstrated consistently higher number of plasma proteins that were altered across all time points. Many upregulated plasma proteins (~303-518) were identified in (+)LC across 5 timepoints, whereas approximately 28-223 proteins were upregulated in (−)LC (Fig 2G). Finally, the metabolomics profiles of COVID19 patients exhibited pronounced alterations in (+)LC patients in most time points in stark contrast to the subtle metabolomics changes in (−)LC group (Fig. 2H and 2I). (+)LC patients showed marked metabolomic alterations at the early timepoint of >-8 days post-COVID19 illness onset compared to (−)LC patients, exhibiting 291 and 6 upregulated metabolites in (+)LC and (−)LC patients, respectively (Fig. 2J). Together, these multi-omics analyses indicate a distinctive immuno-metabolic dysregulation in COVID ‘long haulers’ with chronic headache.

### Hyperinflammation as immunological hallmarks prior to long COVID19 headache onset

To characterize the immune responses that may trigger the onset of long COVID19 headache, we performed bulk RNAseq analysis of blood leukocytes from healthy, (−)LC, and (+)LC groups. Based on the transcriptomic signatures that were differentially altered in (−)LC and (+)LC groups (Fig. 2D), we evaluated the canonical signaling pathways of both COVID19 groups over the course of >-8 days to >3 months post COVID19 illness onset. While (−)LC and (+)LC COVID19 groups exhibited a downward trend of many canonical pathways, (+)LC group exhibited more pronounced downregulation of canonical pathways prior to the onset of headache symptom (>-8 days) (Fig. 3A). Specifically, (−)LC group presented a consistent trend of inhibition of canonical pathways throughout the period of post-COVID19 phase (Fig. 3B). In contrast, (+)LC group demonstrated a predominant activation of canonical pathways from >-8 days up to 30 days post COVID19 illness onset, which was subsequently reversed into the inhibition of canonical pathways at 1-3 months (Fig. 3B). These contrasting observations suggested that (+)LC group was driven by a unique cascade of activated immune signaling pathways. Indeed, 38 canonical pathways were activated in (+)LC group prior to the onset of chronic headache (>-8 days): a majority of those activated canonical pathways were associated with inflammation or immune activation pathways – toll-like receptor (TLR), pyroptosis, chemokine (C-X-C motif) ligand (CXCL) 8, triggering receptor expressed on myeloid cells (TREM) 1, interferon (IFN), interleukin (IL)-6 and nitric oxide synthase, and inducible (iNOS) (Fig. 3C). In contrast, (−)LC group demonstrated the inhibition of inflammatory pathways including IL17F, HMGB1, Th1, IFN and NF-κB pathways (Table S2). Furthermore, (+)LC groups showed markedly increased expression of inflammatory immune mediator genes such as C-type lectin domain Family 4 member E (CLEC4E), TLR-4/8/2/1/10, caspase (CASP) 1, TREM1, N-Myc and signal transducer and activator of transcription (STAT) Interactor (NMI), CXCL8, C-X-C motif chemokine receptor (CXCR) 1/2, CXCL1, S100 calcium binding protein (S100) A8 and S100A9 (Fig. 3D).

**Fig. 3.**
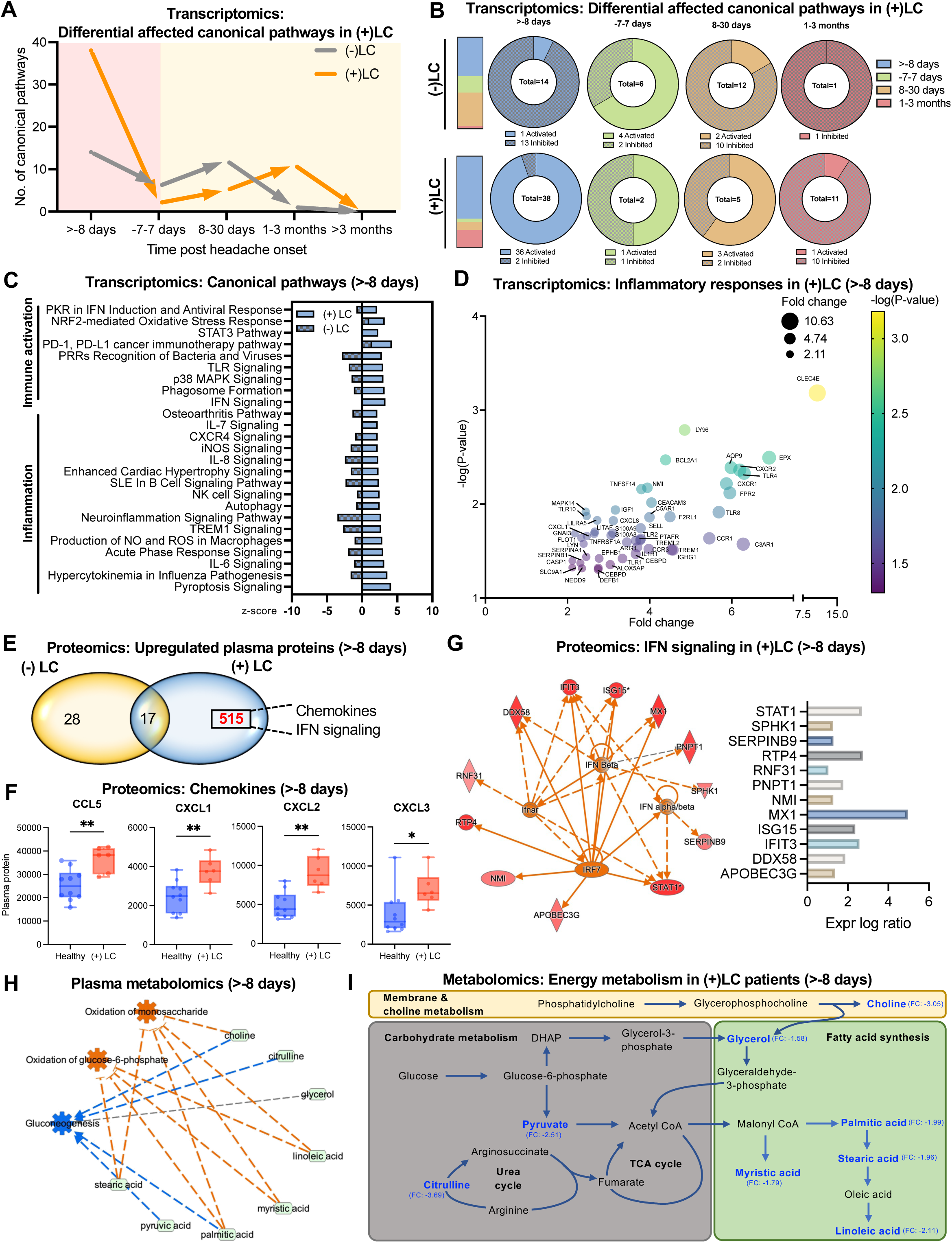
Hyper-inflammation in long COVID patients during early convalescence prior to headache onset. (A-B) Kinetics of the no. of canonical pathways (−2≤ *z*-score ≥2) uniquely affected in (−)LC and (+)LC groups over 5 time points based on transcriptomics profiling. (C) Canonical pathways affected in (−)LC and (+)LC groups at the >-8 days timepoint based on transcriptomics profiling. (D) Bubble plot of transcriptomic expression of inflammatory genes significantly induced in (+)LC group relative to healthy controls at the >-8 days timepoint. (E) Comparison analysis of significantly altered plasma proteins (P < 0.05) present in (−)LC and (+)LC groups at >-8 days timepoint. (F) Chemokines significantly elevated in (+)LC group at >-8 days timepoint. Data are presented as mean ± SEM, using Welch *t*-test. \**P* < 0.05, \*\**P* < 0.01. (G) IFN signaling-associated plasma protein expressions in (+)LC group at >-8 days timepoint. (H) Predicted metabolic pathway activation/inhibition based on plasma metabolomics of (+)LC group at >-8 days timepoint. (I) Overview of energy metabolism in (+)LC group based on plasma metabolomics of (+)LC group at >-8 days timepoint.

Comparative proteomics analysis of plasma specimens identified significantly higher numbers of upregulated plasma proteins such as chemokines and IFN signaling proteins in (+)LC groups (515 proteins or 127 predicted pathways affected) compared to (−)LC groups (28 proteins or 1 predicted pathway affected) (Fig. 3E and S1AB). Those chemokines were CC chemokine ligand 5 (CCL5), chemokine (C-X-C motif) ligand 1 (CXCL1), CXCL2 and CXCL3 (Fig. 3F); and those IFN signaling proteins were STAT1, NMI, MX Dynamin Like GTPase 1 (MX1), IFN-stimulated gene (ISG) 15, IFN-induced protein with tetratricopeptide repeats 3 (IFIT3), DExD/H-box helicase (DDX58, RIG-I) (Fig. 3G).

Ingenuity pathway analysis (IPA) of the metabolomics profiling of (+)LC groups relative to healthy controls demonstrated an activation of glucose metabolism pathways – oxidation of monosaccharide and glucose-6-phosphate – and an inhibition of gluconeogenesis (Fig. 3H). This was consistent with the reduced levels of plasma metabolites involved in the carbohydrate metabolism (pyruvate and citrulline), fatty acid synthesis (glycerol, myristic acid, palmitic acid, stearic acid, and linoleic acid) and membrane/choline metabolism (choline) (Fig. 3I). In summary, multi-omics analysis of blood specimens suggests the hyperactivation of inflammatory responses of (+)LC groups prior to the onset of long COVID19 headache, compared to (−)LC groups.

### Persistent inflammation throughout the long COVID chronic headache progression

To characterize the longitudinal immune landscape of long COVID groups, we performed plasma proteomics analyses over the course of long COVID headache progression. IPA analysis of plasma proteomics profiles of (+)LC groups predicted sustained inflammatory signaling pathways throughout 5 time points post headache onset in addition to hyperinflammatory responses prior to headache onset, which was not detected in (−)LC group (Fig. 4A). Specifically, IFNγ, IFN regulatory factor (IRF) 7, and peroxisome proliferator activated receptor gamma (PPARG) pathways were predicted to be highly activated (Fig. 4A). Indeed, the significantly increased plasma levels of IFNγ and IFNγ-associated downstream cytokines were detected in (+)LC groups compared to healthy controls (Fig. 4B and 4C). IL-6, Immunoglobulin D (IgD), CASP4, and CASP10 were also highly expressed in (+)LC groups, but not in (−)LC groups (Fig. S1B).

**Fig. 4.**
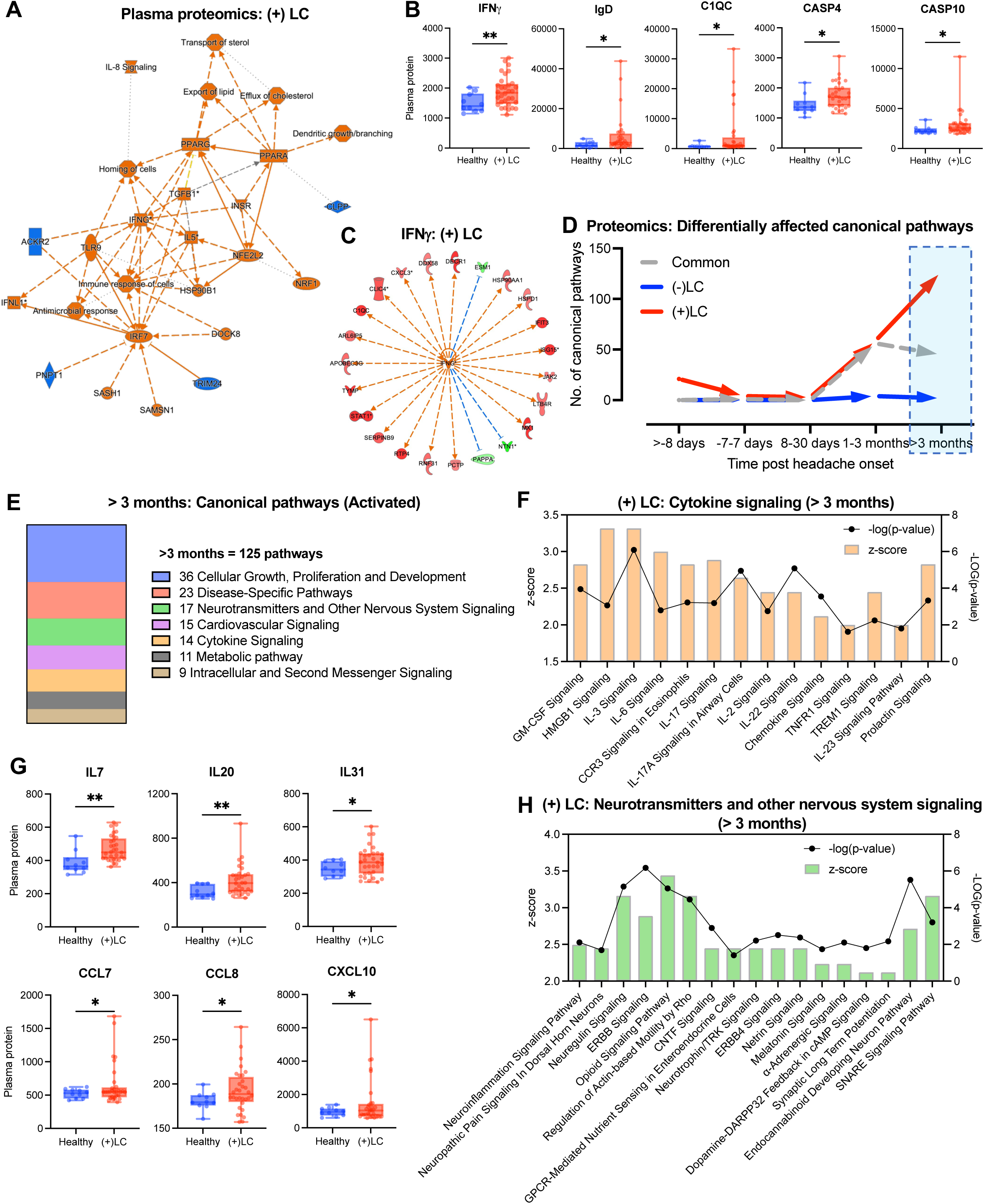
Sustained inflammation in long COVID patients with chronic headache during late convalescence. (A) Graphical summary of immune signaling affected in (+)LC group based on plasma proteomics profiles. (B) Inflammatory immune mediators significantly affected in (+)LC group. (C) Predicted upstream regulator of (+)LC group based on plasma proteomics profiling. (D) Kinetics of the no. of canonical pathways (−2≤ *z*-score ≥2) uniquely affected in (−)LC or (+)LC group over 5 time points based on plasma proteomics profiling. (E) Categorization of canonical pathways that were predicted to be uniquely activated in (+)LC group at >3 months timepoint. (F) Canonical pathways associated with cytokine signaling that were predicted to be uniquely activated in (+)LC group at >3 months timepoint. (G) Cytokines and chemokines elevated in plasma at >3 months timepoint. (H) Canonical pathways associated with neurotransmitters and other nervous system signaling that were predicted to be uniquely activated in (+)LC group at >3 months timepoint. For (B) and (G), data are presented as mean ± SEM, using Welch *t*-test. \**P* < 0.05.

To delineate the plasma proteomics that may contribute to the persistence of chronic headache symptom in (+)LC patients, we performed in-depth analysis of protein biomarkers that were significantly altered in (−)LC and (+)LC groups relative to healthy controls across all time points. Pathway analysis of (−)LC and (+)LC groups were evaluated for (i) common pathways that were altered in both (−)LC and (+)LC groups, and (ii) unique pathways that were altered exclusively in either (−)LC or (+)LC groups (Fig. 4D). While several common pathways were progressively affected in both (−)LC and (+)LC groups over time, a number of pathways were exclusively altered in (+)LC patients, particularly at >3 months post symptom onset (Fig. 4D). Intriguingly, 125 pathways were predicted to be uniquely activated in (+)LC groups at >3 months post headache onset (Fig. 4D). Those pathways are: (i) 36 cell growth, proliferation and development, (ii) 23 disease-specific pathways, (iii) 17 neurotransmitters and other nervous signaling, (iv) 15 cardiovascular signaling, (v) 14 cytokine signaling, (vi) 11 metabolic signaling and (vii) 9 intracellular and second messenger signaling (Fig. 4E).

Inflammation and dysregulation of neurotransmitters have been associated with the development of various forms of migraine, episodic and tension headaches. 14 cytokine signaling and 17 neurotransmitters signaling pathways were specifically altered in (+)LC groups, but not (−)LC groups (Fig. 4F–4H). Activated cytokine signaling pathways were associated with inflammation such as GM-CSF, IL-6, IL-17, IL-2, IL-22, TNFR1, IL-23, chemokines, etc. (Fig. 4F). Indeed, elevated plasma levels of inflammatory cytokines including IL-7, IL-20, IL-31, CCL7, CCL8 and CXCL10 were clearly detected in (+)LC groups compared to healthy controls (Fig. 4G). Additionally, 17 activated neurotransmitter signaling pathways included neuroinflammation, neuregulin signaling, opioid signaling, melatonin signaling, dopamine signaling, endocannabinoid pathways, and others. (Fig. 4H). This longitudinal analysis of plasma proteomics profile reveals a sustained inflammatory response and dysregulated neuro-signaling pathway that may contribute to the progression of chronic headache in (+)LC groups.

### Immuno-metabolic regulation in long COVID headache patients

Metabolism plays a crucial role in regulating inflammation often through mediating immune cells activation and differentiation^25^. LC-MS-based metabolomics analysis of the longitudinal plasma specimens of healthy and COVID groups revealed suppression of citrulline biosynthesis/metabolism and arginine degradation in (+)LC groups, suggesting a dysregulation of urea cycle (Fig. 5A). Specifically, Arginine, a key metabolite in urea cycle, was one of the top metabolites that was upregulated in (+)LC groups, but not in (−)LC groups (Fig. 5B). Individual timepoint analysis further revealed the upregulated levels of arginine as early as −7-7 days up till >3 months post illness onset (Fig. 5C). Similarly, oxo-arginine, a metabolite of arginine catabolism, significantly increased at the later phase of long COVID illness, (Fig. 5C).

**Fig. 5.**
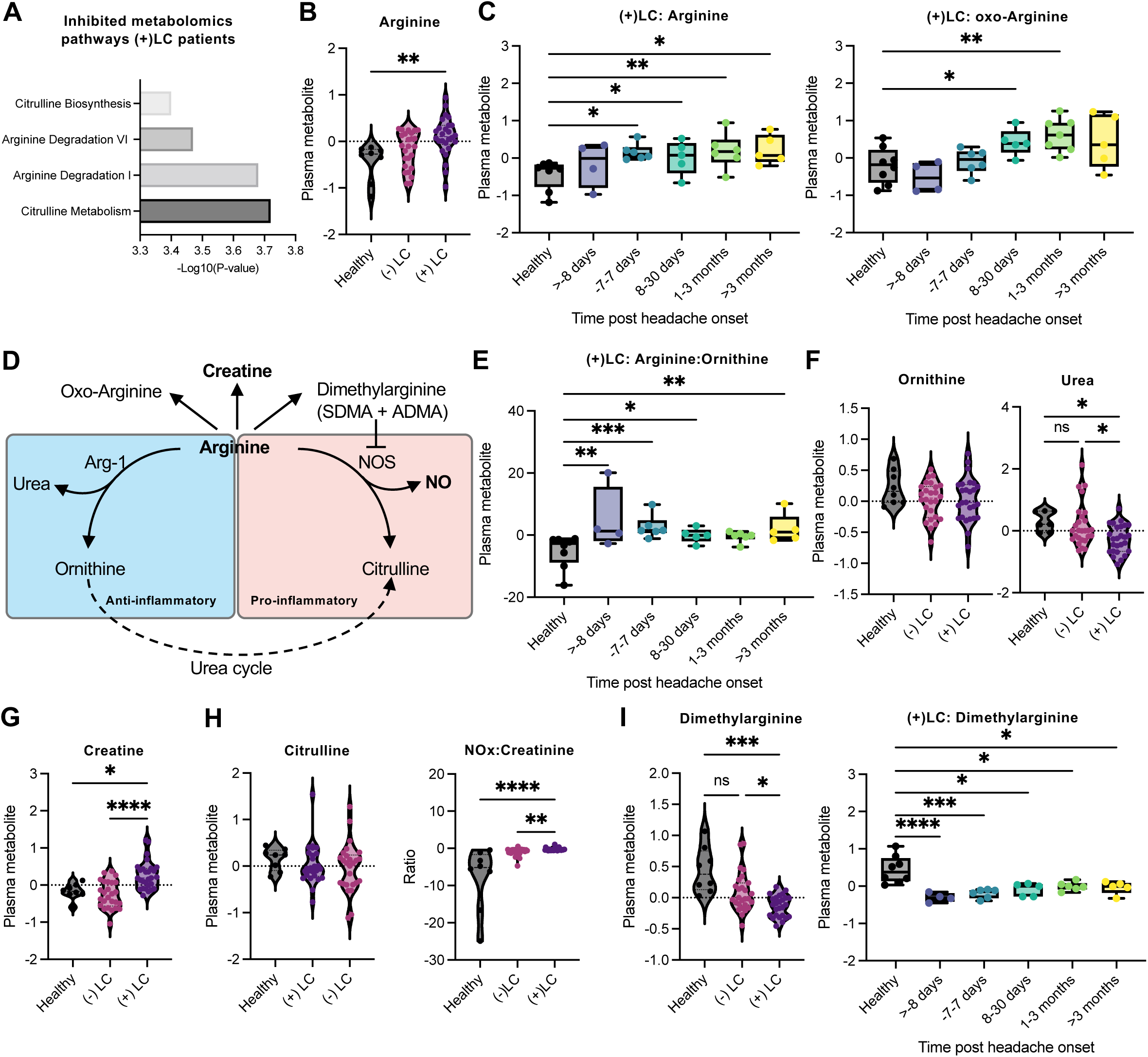
Immuno-metabolic modulation of hyperinflammation during long COVID associated chronic headache. (A) Metabolic pathways predicted to be inhibited in (+)LC group based on plasma metabolomics profiling. (B-C) Plasma levels of arginine and arginine-derivative, oxo-Arg. (D) Plasma metabolites associated with the Urea cycle. (E-F) Plasma levels of metabolites associated with anti-inflammatory arm of arginine metabolism – Arg-Orn ratio (E), Orn and Urea (F). For (G-I), plasma levels of metabolites associated with pro-inflammatory arm of arginine metabolism – Creatine, Cit, NO, dimethylarginine. Data are presented as mean ± SEM, using One-way ANOVA Kruskal-Wallis with uncorrected Dunn’s test. \**P* < 0.05, \*\**P* < 0.01, \*\*\**P* < 0.001, \*\*\*\**P* < 0.0001.

Arginine has been shown to play a central role in modulating macrophage polarization through either arginase-1-dependent anti-inflammation or nitric oxide synthase (NOS)-dependent pro-inflammation^26^. Arginine is converted to (i) ornithine and urea, (ii) citrulline and NO or (iii) creatine (Fig. 5D). Abundance of plasma arginine was clearly evident when arginine:ornithine levels were significantly elevated as early as >8 days prior to headache onset, up to > 3 months after headache onset (Fig. 5E), along with a significant reduced level of urea in the (+)LC patients (Fig. 5F). However, neither ornithine nor citrulline level was significantly altered in all COVID groups (Fig. 5F and 5H), whereas creatine level was elevated in (+)LC groups (Fig. 5G). As NO is rapidly oxidized to nitrates and nitrites (NOx) by oxyhemoglobin, NOx levels were measured to assess plasma NO production. In addition, since both NOx and creatinine are excreted renally, the plasma NOx:creatinine ratio can be used as an indicator of the endogenous NO production^27,28^. While plasma NOx and creatinine levels were reduced in both (−) LC and (+)LC patients compared to healthy controls (Fig. S2), the NOx/creatinine ratios were detectably elevated in (+)LC patients when compared to (−)LC patients and healthy controls (Fig. 5H and Fig. S2). Furthermore, levels of asymmetric and symmetric dimethylarginine (ADMA and SDMA), which are potent inhibitor of NOS activity^29^, was significantly reduced specifically in (+)LC group (Fig. 5I). These data highlight a potential role of skewed Arginine-NO metabolic pathway in sustained inflammation among the (+)LC headache groups.

### Increase of lipid metabolism during long COVID headache progression

Inflammation is an energy-consuming mechanism of host defense. So, it partly relies on lipid metabolism to facilitate macrophage differentiation and function^30^. Lipid metabolism can fuel both pro-inflammatory M1 and anti-inflammatory M2 macrophage differentiation through fatty acid synthesis and fatty acid oxidation metabolic pathways, respectively. Hence, abundance of free fatty acids (FFAs) has been linked to pro-inflammatory macrophage activation^30^. When we compared the significantly upregulated metabolites (relative to healthy controls) between (−)LC (10 metabolites) and (+)LC (56 metabolites) groups, 51 metabolites were uniquely upregulated only in (+)LC group (Fig. 6A). Metabolic pathway analyses revealed that a majority (~67%) of 51 (+)LC upregulated metabolites were from lipid-dominating super pathways (Fig. 6B). Specifically, the top 5 upregulated metabolic pathways were (i) sphingolipid metabolism, (ii) glycerophospholipid metabolism, (iii) phenylalanine, tyrosine, and tryptophan biosynthesis, (iv) linoleic acid metabolism, and (v) arginine and proline metabolism (Fig. 6C). Highly upregulated lipid metabolites were 16 sphingomyelin derivatives, 7 plasmalogen lipid derivatives, 3 phospholipid derivatives, 3 sphingolipid derivatives, 3 glycosphingolipid derivatives and cholesterol (Fig. 6D). Levels of all 34 lipid metabolites were considerably higher in (+)LC groups compared to (−)LC groups. Specifically, sphingomyelin derivatives (d18:2/24:1, d18:1/24:2) and (d18:2/16:0; d18:1/16:1) (Fig. 6E), plasmalogen lipids — 1-(1-Enyl-palmitoyl)-2-oleoly-GPE (P-16:0/18:1) and 1-(1-Enyl-oleoyl)-GPE (P-18:1) (Fig. 6F), phospholipid — 1-stearoyl-2-oleoyl-GPS (18:0/18:1) (Fig. 6G), sphinganine (Fig. 6H), glycosyl ceramide (d18:1/20:0, d16:1/22:0) (Fig. 6I) and cholesterol (Fig. 6J) were significantly elevated throughout time points of headache progression in (+)LC patients. Herein, our data presented an elevated lipidomics profile in (+)LC patients, implicative of a FFAs-driven chronic inflammation in conjunction with headache progression.

**Fig. 6.**
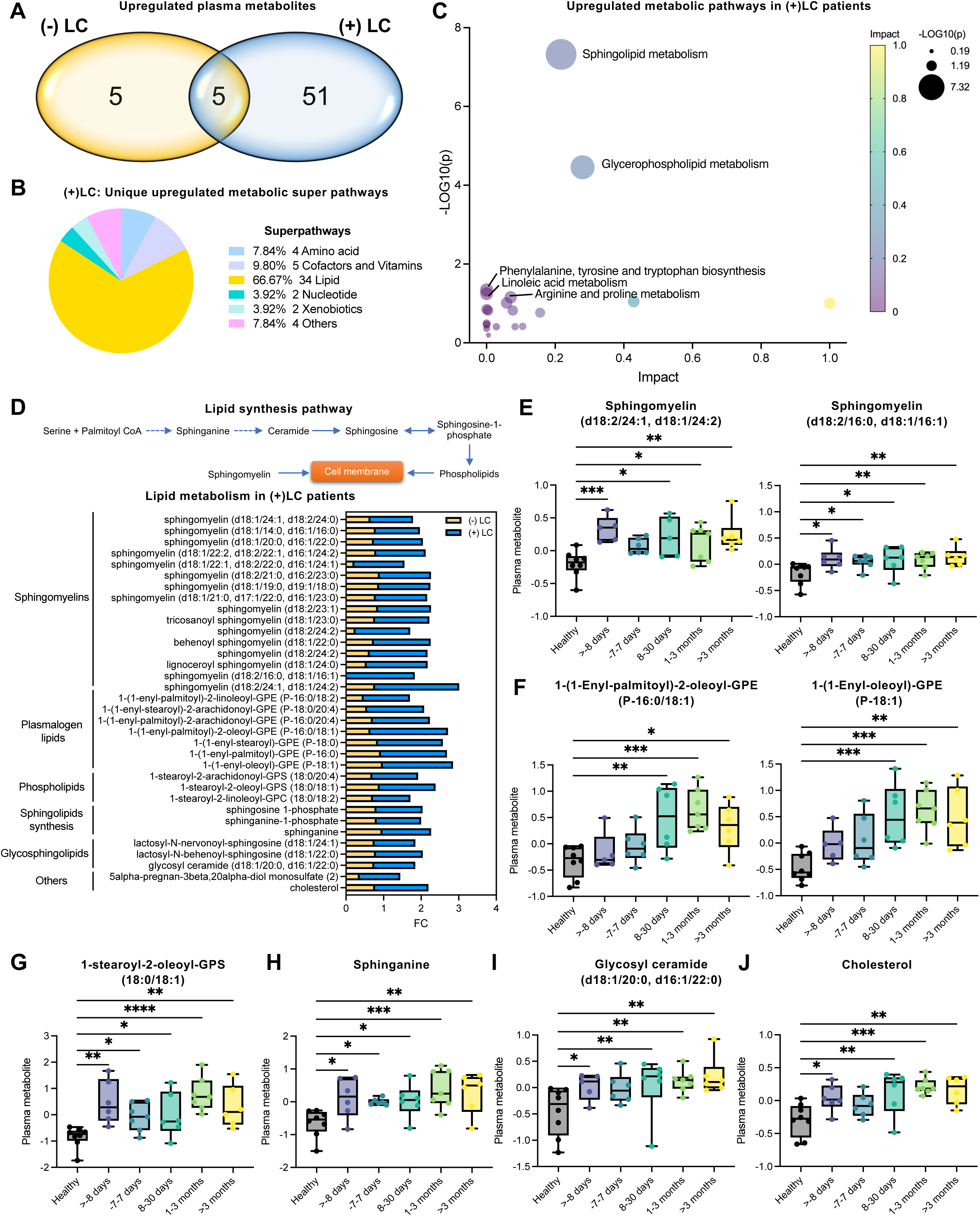
Augmented phospholipid metabolism in long COVID-headache patients. (A) Comparative analysis of significantly altered plasma metabolites (P < 0.05) present in (−)LC and (+)LC groups. (B) Categorization of upregulated metabolic super-pathways in (+)LC group. (C) MetaboloAnalyst prediction of upregulated metabolic pathways in (+)LC patients. (D) Lipid metabolism-associated plasma metabolites significantly altered in (+)LC patients. (E-J) Levels of plasma metabolites associated with lipid metabolism in (+)LC group: specifically sphingomyelins (E), plasmalogen lipids (F), phospholipids (G), sphingolipids synthesis (H), glycosphingolipids (I) and cholesterol (J). Data are presented as mean ± SEM, using One-way ANOVA Kruskal-Wallis with uncorrected Dunn’s test. *P < 0.05, **P < 0.01, ***P < 0.001, ****P < 0.0001.

### Neurotransmitters and neuroactive metabolites dysregulation during long COVID headache

Unbalanced levels of neurotransmitters such as tryptophan, serotonin (tryptophan-derived), tyrosine, dopamine (tyrosine-derived), glutamate and GABA (glutamate-derived) have long been associated with headaches^31^. Pathway analysis of both plasma proteomics and metabolomics pinpointed towards a dysregulation of neurotransmitters signaling and tyrosine/tryptophan biosynthesis in (+)LC patients (Figs. 4H and 6C). Interestingly, while tryptophan level was not altered in both COVID19 groups, kyneurenine level was decreased and serotonin level was increased in (+)LC groups when compared to (−)LC or healthy groups (Fig. 7A). In fact, sustained increases of serotonin levels were observed throughout the timepoints in (+)LC groups (Fig. 7B). Increased serotonin level could affect dopamine level through the competition for aromatic L-amino acid decarboxylase (AADC) enzyme which can convert levodopa (L-DOPA) into dopamine or 5-hydroxytryptophan (5-HTP) into serotonin^32^ (Fig. 7C). The levels of tyrosine and ADCC were similar across all three groups (Fig. 7D and 7E). When compared to (−)LC patients or healthy controls, the levels of dopamine intermediates, dopamine-3-O-sulfate and dopamine-4-O-sulfate, were significantly declined in the (+)LC group at the early phase of long COVID19 illness between >-8 days to 7 days post headache onset (Fig. 7F and 7G). α-ketoglutarate serves as the precursor for the synthesis of glutamate and γ-aminobutyric acid (GABA), which are both neurotransmitters^33^. The reduced levels of α-ketoglutarate, glutamate and carboxyethyl-GABA (GABA derivative) were exclusively detected in (+)LC groups in almost all time points when compared to healthy groups (Figs. 7H-7J). The reduced GABA level in (+)LC groups was further evidenced by plasma proteomics profiling analysis that predicted GABA inhibition along with induced levels of downstream proteins (Fig. 7K). In summary, our data demonstrate the dysregulation of multiple neurotransmitters throughout the progression of chronic headache in (+)LC groups.

**Fig. 7.**
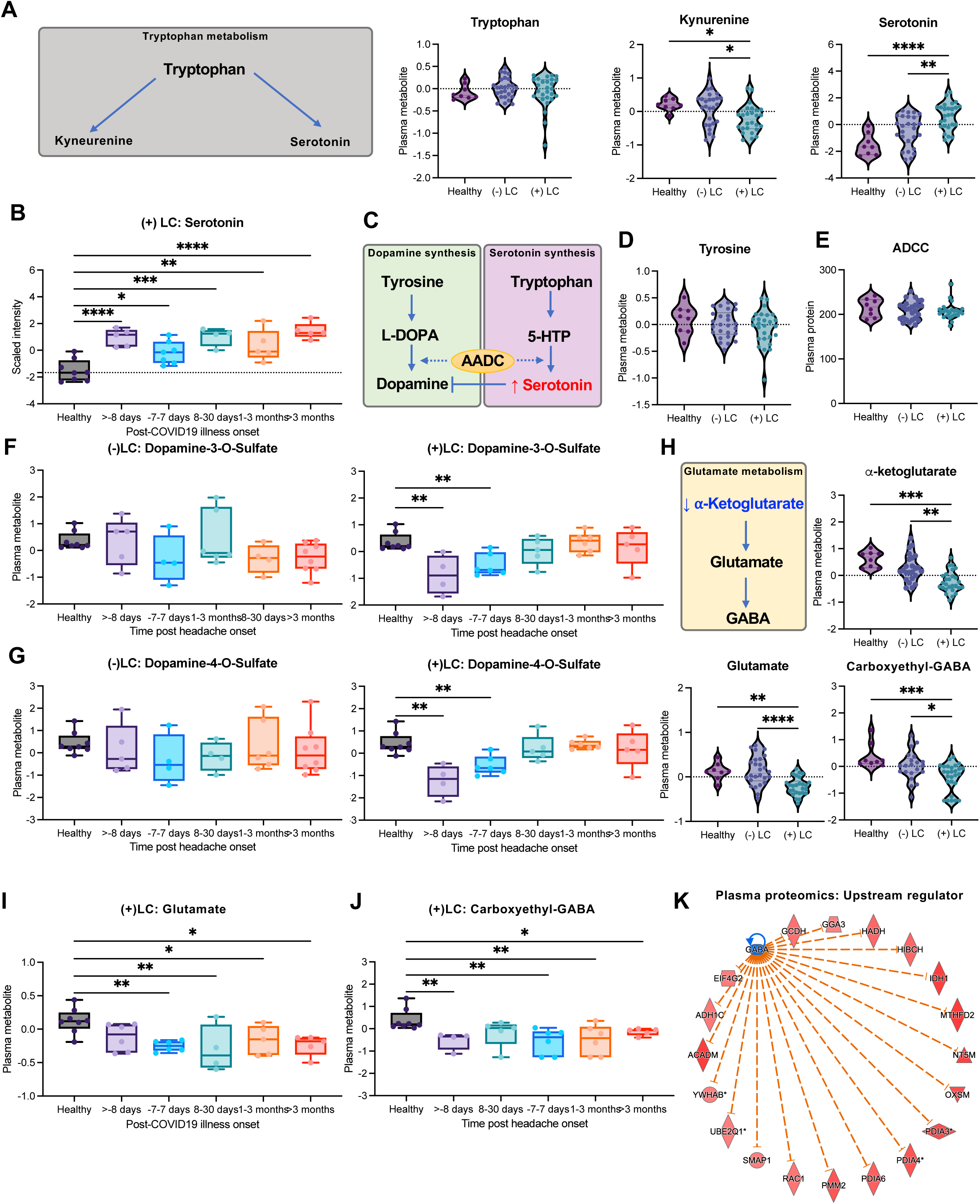
Dysregulated neurotransmitters and neuroactive metabolites in long COVID-headache patients. (A) Levels of plasma metabolites associated with tryptophan metabolism. (B) Serotonin level in (+)LC patients’ plasma specimens. (C) Schematic overview of dopamine and serotonin metabolic synthesis pathways. (D-E) Levels of plasma metabolites/protein associated with dopamine synthesis pathway. (F-G) Dopamine metabolites in (−)LC and (+)LC patients’ plasma specimens. (H) Levels of plasma metabolites associated with glutamate metabolism. Glutamate (I) and carboxylethyl-GABA (J) levels in plasma specimens of (+)LC patients. (K) GABA as upstream regulator of plasma proteomics profiles from (+)LC group. For (A, B, D-J), data are presented as mean ± SEM, using One-way ANOVA Kruskal-Wallis with uncorrected Dunn’s test. \**P* < 0.05, \*\**P* < 0.01, \*\*\**P* < 0.001, \*\*\*\**P* < 0.0001.

## DISCUSSION

The prevalence of post-COVID symptoms in COVID-19 survivors is an emerging global health concern. Post-COVID symptoms can include a wide spectrum of respiratory, heart digestive and neurological problems at ≥12 weeks after infection that can last for months or years^34^. In fact, neurological symptoms such as fatigue and chronic headaches are among the most common symptoms presented by long COVID patients. A recent cohort study has reported over 66.5% (133/200) of COVID-19 patients suffering from lingering headaches about 4 months after testing positive for the virus^35^ and yet, the etiology of long COVID-related chronic headaches is still not well characterized. Here, we used retrospective longitudinal blood specimens from early to late convalescence COVID-19 patients (prior and after the first headache diagnosis) experiencing prolonged headache and healthy controls to characterize the immuno-metabolic landscape in the onset of chronic headaches during acute infection phase and convalescence phase of PASC. This study illustrates a novel immuno-metabolomics landscape of long COVID patient with chronic headaches that may provide insights to future therapeutic interventions.

It is well established that elevated C-reactive protein (CRP) is linked to the pathogenesis of migraines and associated with a low pain threshold in headaches^36,37^. In a cross-sectional study of headache attributed to SARS-CoV-2 infection, CRP levels have been reported to be elevated without specific association with severity and frequency of COVID-19 related headache^38,39^. We also observed higher CRP levels in COVID-19 patients compared to healthy controls at the time of positive COVID-19 diagnosis (Table S2). However, the CRP levels of both (−)LC patients and (+)LC patients fell to the moderately elevated ranges (1-10mg/dL), suggesting that CRP level may not be a right inflammation predictor of long COVID headaches in the early and late convalescence phase of COVID-19. As previously described for pain intensity^38^, dehydration was also present in our (+)LC patients, suggesting a potential predictor of long COVID chronic headache. Although CBC tests of our (+)LC patients were within normal ranges, (+)LC patients showed modest increases of RBC, hemoglobin and hematocrit levels at early convalescence and detectable increases of platelet and WBC counts at late convalescence compared to (−)LC patients. These suggests that there are several potential phenotypes associated with (+)LC patients with chronic headache.

Our COVID19 patient cohorts regardless of chronic headache either were asymptomatic or displayed mild COVID19 disease. Despite seemingly mild COVID19 illness during the acute stage of infection, many of these patients progressed into long COVID headache during the convalescence phase of infection. We found that the onset of long COVID headache might be triggered by a state of hyperinflammation which was sustained throughout the progression of chronic headache even months post infection. Neuro-inflammation has been well associated with the pathophysiology of migraine-like headache attacks^40–42^. Indeed, recent single-nucleus transcriptomic study of brain tissues of deceased severe COVID19 patients revealed the evidence of neuroinflammation involving the upregulation of IFN, chemokine and complement pathways^43^. Consistent with this study, our transcriptomics and proteomics analyses also detected elevated levels of IFN-associated factors (*e.g.,* STAT1, ISG15, IFIT3), chemokines (*e.g.,* CCL5, CXCL1/2/3/10), and other inflammatory cytokines such as IFNγ, IL-6, IL-7/20/31. These observations imply that chronic systemic inflammation may drive the development of long COVID headache regardless of disease severity.

Immunometabolism is emerging as a crucial network of sophisticated intertwined metabolic processes which regulate immune activation through the generation of amino acids, metabolites, neurotransmitters and energy^44^. Glycolysis is one of the primary metabolic pathways in driving energy production to fuel the downstream cascade of metabolic processes including tricarboxylic acid (TCA) and urea cycles^45^, as well as energy-consuming proinflammation^46^. Herein, we observed an increased glucose metabolism in long COVID headache patients in conjunction with a myriad of activated inflammatory responses prior to the onset of headache symptom, which potentially served as an energy generation source to ignite the onset of hyper-inflammation. Intriguingly, despite viral clearance, long COVID patients continued to sustain low-grade inflammation, evidenced by persistent increases of inflammatory markers such as IL-6 and IFN-γ and chemokines such as CXCL10, CCL7 and CCL8 even up to more than 3 months post headache symptom onset.

Chronic low-grade metabolic inflammation, also known as “metainflammation”, is the hallmark of metabolic disorders including obesity, type 2 diabetes, non-alcoholic fatty liver disease and old age-associated chronic inflammation (inflammaging)^47–49^. Dysregulation of lipid metabolism has been a key driver of metainflammation during obesity, in which increase levels of lipids such as triglycerides (TAG), diacyglycerol (DAG), sphingolipids (sphingomyelins, ceramide, sphingosine and sphingosine-1-phosphate) in the circulation can lead to elevated proinflammatory cytokines production, systemic inflammation and lipotoxicity^48,50^. Remarkably, numerous lipid metabolites, including sphingomyelins, plasmalogen, phospholipids, sphingolipids, and cholesterol, were elevated exclusively in the plasmas of long COVID patients throughout the progression of chronic headache. This phenomenon not only recapitulated the lipotoxicity observed in metabolic disorders such as obesity, but also highlighted the implication of a dysregulated lipid metabolism underpinning meta-inflammation-like conditions in long COVID-headache patients.

Metabolic reprogramming through lipid metabolism has been well associated with the polarization of macrophages^51^. Inflammatory M1 macrophages undergo lipid biosynthesis to facilitate membrane remodeling and production of inflammatory lipids, distinct from the fatty acid oxidation process that anti-inflammatory M2 macrophages require^51^. Lipidomic profiling of M1 macrophages revealed the increased production of glycerolipids, glycerophospholipids and sphingolipids that are essential for triggering downstream inflammatory signaling^48,52^. Notably, these lipid metabolites were also markedly elevated in long COVID patients, indicative of M1 macrophage-induced inflammation. Apart from lipid metabolism, arginine metabolism also plays an indispensable role in macrophage polarization, in which pro-inflammatory M1 macrophages convert arginine to citrulline and nitric oxide (NO) through NO synthase (NOS), while anti-inflammatory M2 macrophages convert arginine to ornithine and urea through arginase-1^53^. Besides M1-skewed lipidomic profiles observed in long COVID-headache patients, we also identified the increased levels of arginine and its metabolite oxo-arginine along with NO, which were accompanied by the reduced levels of urea and NOS inhibitors (SDMA and ADMA). These findings further supported the critical roles of proinflammatory M1 macrophage-driven sustained inflammation in long COVID, likely contributing to chronic headache. NO, a small gaseous signaling molecule, has been heavily implicated in various forms of headaches including migraine or cluster and tension headaches due to its vasodilation and pain processing properties^54^. As a byproduct of the arginine-citrulline biochemical pathway in M1 macrophages, NO levels were augmented in long COVID patients, fueling chronic headache symptom. Hence, NOS inhibitor drugs that curb the production of NO may be a potential therapeutic strategy for relieving persistent inflammation and headache in long COVID patients.

The etiology underlying the pathophysiology of headaches have been an open debate, especially in the aspect of chronic headache. Chronic headaches are distinct from episodic headaches like migraine, which involves a sustained transmission of pain sensation following the trigger of headache onset. Neurotransmitters such as glutamate, GABA, dopamine and serotonin are produced biochemically through tyrosine, tryptophan or glutamate metabolism, respectively, which can mediate headache onset and modulate pain sensation^55^. Fluctuation of glutamate-derived GABA levels has been associated with the severity of headache attacks, whereby increased GABA dampens pain tolerance^56,57^. For instance, while increased levels of GABA have been reported during tension headache^58^ as well as during migraine attacks compared to headache-free periods^59^, reduced GABA levels have been also observed in patients with moderate/severe migraine at 1 month after headache onset^57^. Similarly, reduced levels of dopamine can also contribute to the development of chronic pain^60^. Abnormalities in serotonin metabolism have been reported during headache attacks: specifically, low levels of serotonin were detected in patients suffering from tension-type headache^61^. Our long COVID headache patient cohort also showed dysregulation of these neurotransmitters. Specifically, serotonin was significantly elevated across all timepoints even before headache onset, whereas dopamine metabolites, glutamate and GABA metabolites were significantly reduced during the same periods. Together, our data suggest critical roles of dysregulated neurotransmitter metabolisms in the progression to chronic headache in long COVID patients.

In conclusion, using multi-omics approach, we dissected the immuno-metabolic profiles of a unique cohort of COVID patients who developed long COVID-associated chronic headache symptom. Specifically, we identified that: (i) hyperinflammation was involved in triggering the onset of long COVID-headache symptom, (ii) sustained inflammation might contribute to the development of long-term post-COVID headache, (iii) immuno-metabolic reprogramming in long COVID-headache patients might drive the accumulation of arginine and lipid metabolites to fuel sustained inflammation, and (iv) dysregulated metabolism of multiple neurotransmitters might be the hallmark of long COVID chronic headache (Fig. 8). Together, our findings shed light on understanding the progression of chronic headache of long COVID patients and open the development of targeted therapeutic strategies for treatment of this debilitating condition.

**Fig. 8.**
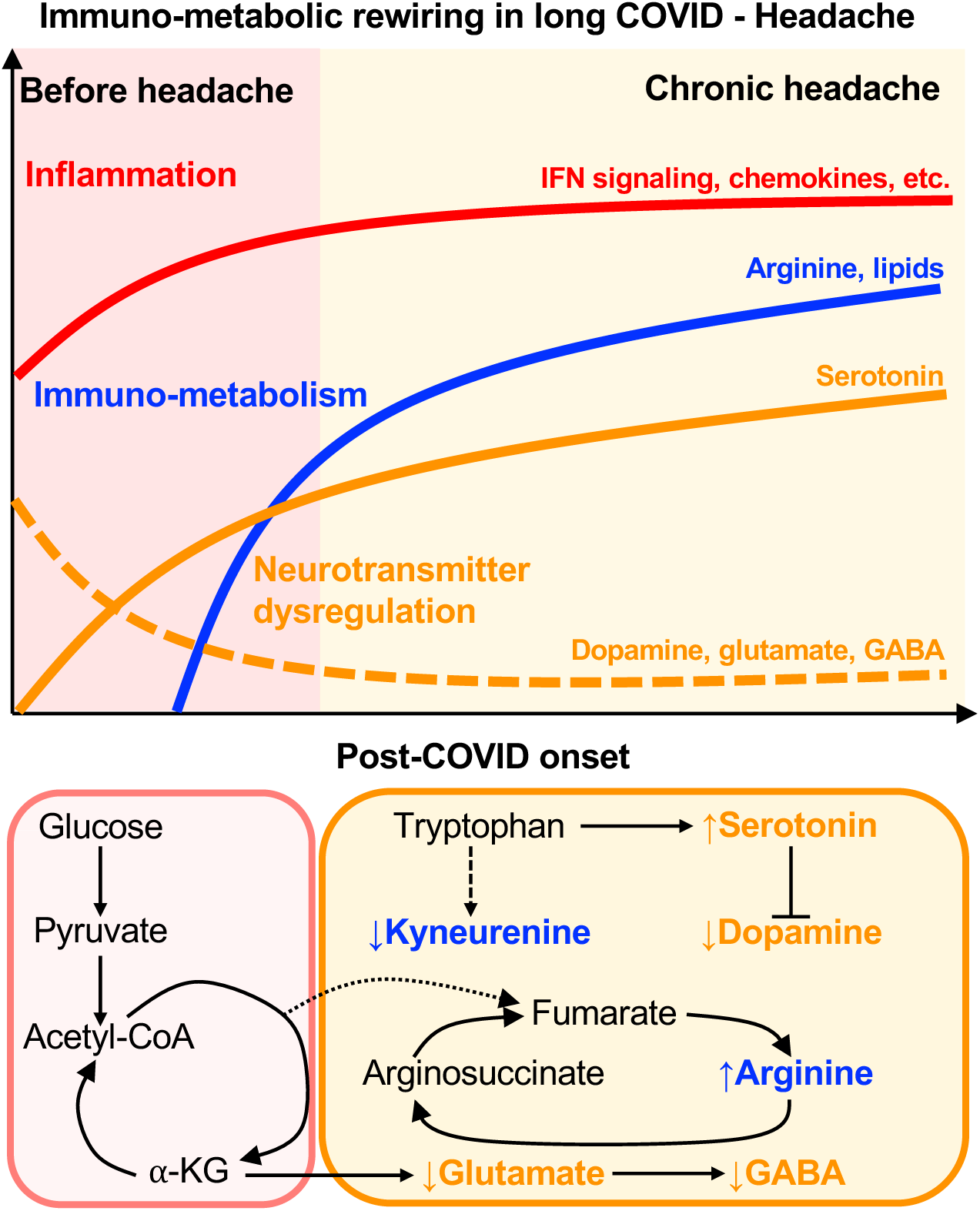
Overview of immuno-metabolic rewiring in long COVID patients with chronic headache.

## MATERIALS AND METHODS

### COVID19 patient cohort and blood specimen collection

Herein, a retrospective clinical cohort of adult COVID19 patients (>18 years of age) with qRT-PCR confirmed SARS-CoV-2-positive nasopharyngeal swab test were enrolled for this long COVID study. Clinical testings including SARS-CoV-2 qRT-PCR testing and complete blood count were performed at the Cleveland Clinic Pathology & Laboratory Medicine Institute (Robert J. Tomsich) between May 2020 and Oct 2021. The cohort was followed longitudinally after COVID-19 diagnosis with peripheral blood collection at various time points post-acute COVID19 diagnosis. All COVID19 patients (with or without long COVID) were clinically evaluated according to acute-phase symptoms, smoking habits, the existence of co-morbidities, and the baseline medication used at the time of COVID diagnosis. Patients were categorized as long COVID (LC) group if headache was present in the convalescence phase. Clinical phenotyping is systematically done through the Cleveland Clinic COVID Registry, capturing all patients tested for COVID across the Cleveland Clinic health system, now including >500k patients, with>80 datapoints per patient. A detailed description of which has been published previously^62^. The long COVID blood samples were collected between early to late convalescence phase, with early convalescence phase defined as: (i) less than 8 days before (<-8 days) or (ii) 7 days before and after (−7 to 7 days) headache onset; whereas, late convalescence phase are defined as (iv) between 8 and 30 days (8-30 days), (v) 1 to 3 months (1-3 months), or (vi) more than 3 months (>3 months) after headache onset. For the (−)LC patients, blood samples were collected at similar timepoints as outlined above, based on post-SARS-CoV-2 positive testing. Healthy controls were age- and sex-matched to COVID19 patients’ cohort. For all blood specimens collected, packed cells (resuspended in RNALater) and plasma samples were isolated and stored at −80 °C. The study cohort and sample collection are approved by the Cleveland Clinic Institutional Review Board (IRB) and Institutional Biosafety Committee (IBC). Informed consent for study participation was obtained for all participants before enrollment.

### Transcriptomic profiling – RNA Sequencing

Total RNA was extracted from packed blood cells resuspended in RNALater using RiboPure™ RNA Purification (blood) Kit (Invitrogen, USA) as per manufacturer’s protocol. Purified total RNA was processed into RNA sequencing libraries with the KAPA mRNA Hyperprep Kit (Roche, USA). Quality control was tested before and after library preparation using an Agilent TapeStation 4200 and sequencing was performed by the Cleveland Clinic Genomics Core using the Illumina sequencing platform NovaSeq 6000 (Illumina, USA). De-multiplexed fastq files were received from the Genomics Core and utilized for further analysis. Fastq files were analyzed with Partek® Flow® software, version 10.0.21.1026, (Partek, USA). Reads were trimmed and aligned to the human genome hg38 using STAR aligner and gene read counts were obtained using the Partek Expectation/Maximization (E/M) algorithm, transcript model hg38-RefSeq Transcript 94. Genes with average coverage ≥ 1 read per million were defined as expressed and selected for differential gene expression analysis using the Gene set Analysis (GSA) method (lognormal with shrinkage). Significantly differentially expressed genes between the groups analyzed were identified in Partek Flow by fold-change ≥ 2 and FDR-adjusted p-value < 0.05.

### Plasma proteomics profiling

Plasma samples were analyzed using the SomaScan assay (SomaLogic, USA). Briefly, SomaScan assay contains 7000 aptamers that recognize specific protein antigens. After binding, the levels of protein-aptamer complexes are quantified using next-generation sequencing ^63^. Raw data in .adat format was read into R environment with the readat R package ^64^. Fold change was calculated by first averaging the expression of each group and then using the following equation: (COVID-19 LC(−) or COVID-19 LC(+) expression – Healthy expression)/ (Healthy expression). This formula accounts for negative expression values that were generated during the initial log-2 conversion and normalization. Significant differentially expressed proteins were determined by Welch t-tests or Wilcoxon Rank Sum Tests using the R base package, t-test, considering fold-change ≥ 2 and FDR-adjusted p-value < 0.05.

### Plasma metabolomics profiling

The Metabolon company (Morrisville, North Carolina, USA) conducted the metabolomics assays for all participant plasma samples used in this study. The Global Metabolomics platform was used to generate quantitation of 1,400 metabolites by ultra-high-performance liquid chromatography/tandem accurate mass spectrometry as described previously ^24^. Briefly, 150μl of plasma was transported to Metabolon Inc. for analysis. Mass spectrometry was performed using Metabolon’s ultra-high-performance liquid chromatography/tandem mass spectrometry (UHPLC/MS/MS) Global Platform. Data extraction, along with biochemical identification, data curation, quantification, and data normalizations were performed by Metabolon’s hardware and software. Analysis of raw metabolomics data was conducted in R environment ^64^. FC calculation and determination of significantly differentially expressed metabolites were carried out as described in the previous topic.

### Nitric oxide plasma detection

Plasma samples were tested for detection of total Nitric Oxide (NO) using the kit Nitrate/Nitrite Fluorometric Assay kit (Cayman Chemical, USA) following the manufacturer’s instructions. Briefly, 100ul of plasma was de-proteinized using 10kD Spin Columns (Abcam, Cambridge, United Kingdom). Deproteinized samples were analyzed in duplicates in both nitrate + nitrite (total NOx) and nitrite assay. Plates were read at excitation wavelength of 320nm and an emission wavelength of 430nm using the plate reader Varioskan Lux, and data acquisition with the Skanlt microplate reader software (Thermo Scientific, USA). To obtain total NO and nitrite values in µM, the fluorescence of each sample was interpolated with the standard curves using 4 parameters logistic regression, with reduction of blank wells at the Graphpad PRISM 9.0 software (GraphPad Software, San Diego, California, USA). Nitrate concentration was calculated by total NO–nitrite for each sample analyzed.

### Data analysis

Data from RNAseq, proteomics, and metabolomics were submitted to an exploratory analysis conducted in R environment ^65^ or in Graphpad PRISM 9.0 software (GraphPad Software, San Diego, California, USA). tSNE plots were generated in R using the Rtsne package ^66,67^. Enrichment analysis of the canonical pathways of the significantly differentially expressed genes, proteins, or metabolites was conducted using Ingenuity Pathway Analysis (QIAgen Inc., Hilden, Germany). Further statistical tests were done either in R (R Core Team, 2020) or GraphPad PRISM 9.0 software (GraphPad Software, San Diego, California, USA). Comparisons between the two groups were performed using the Welch t-test. For comparisons between more than two groups or time points, one-way ANOVA Kruskal-Wallis with Dunn’s post-test was used.

## Acknowledgments

This work was partly supported by National Institutes of Health (NIH) grants: K99DE028573, R00DE028573 (W.C.), R01NS097719 (L.J.), R35CA200422, R01CA251275, R01AI140705, R01AI140705S, R01AI140718, R01AI152190, R01AI171443, R01AI171201, R01DE023926, R01DE028521 and Korea Research Institute of Bioscience and Biotechnology Research Initiative Program KGM9942011 (J.U.J.).

## Author contributions

S.-S.F., W.C. and J.U.J. designed the experiments. S.-S.F., W.C., K.L.J., T.A. performed the experiments and analyzed the data. S.-S.F. and W.C. prepared the figures and wrote the manuscript. K.L.J., T.A., U.Y.C., P.Z. assisted with the experiments. S.A.A.C., S.C.E., L.J. provided expertise and tools for the study. J.U.J., S.-S.F. and W.C. edited the manuscript and oversaw all study designs and data analyses. All other authors read the manuscript and provided comments.

## Competing interests

Authors declare that they have no competing interests.

## Data and materials availability

The raw data for all graphs generated in this study are provided in the Supplementary Information and Source Data file. Source data are provided with this paper.

## Supplemental information

**Table S1. Overview of the clinical demographics and parameters of COVID19 patients upon COVID19 diagnosis.**

**Table S2. Canonical pathways that are uniquely altered in (−)LC patients.** Canonical pathways with z-score ≥ 2 that were uniquely altered in (−)LC but not in (+)LC group over 5 different time points. **For Fig. 2B**.

**Fig. S1. Immuno-metabolic profiles of (−)LC patients.** (A) Protein biomarkers that are significantly altered in (−)LC or (+)LC compared to healthy controls were subjected to IPA pathway analysis to predict the canonical pathways that were affected in both groups. (B-C) Plasma proteomics profiles of chemokines (B) and inflammatory mediators (C) in (−)LC group. Data are presented as mean ± SEM, using Welch *t*-test. \**P* < 0.05, \*\**P* < 0.01.

**Fig. S2. Nitric oxide biosynthesis in COVID patients.** (A) Nitrite/nitrate and (B) total NOx levels were determined from the plasma samples of both healthy and COVID19 patient groups. (C) Plasma creatinine levels were determined using LC-MS metabolomics profiling. Data are presented as mean ± SEM, using One-way ANOVA Kruskal-Wallis with uncorrected Dunn’s test. \**P* < 0.05, \*\**P* < 0.01, \*\*\**P* < 0.001, \*\*\*\**P* < 0.0001.

## Notes

### Competing Interest Statement

The authors have declared no competing interest.

