## Supplementary figures and images for "Immunometabolic rewiring in long COVID patients with chronic headache"

### Figure S1

**Fig. S1. Immuno-metabolic profiles of (-)LC patients.**

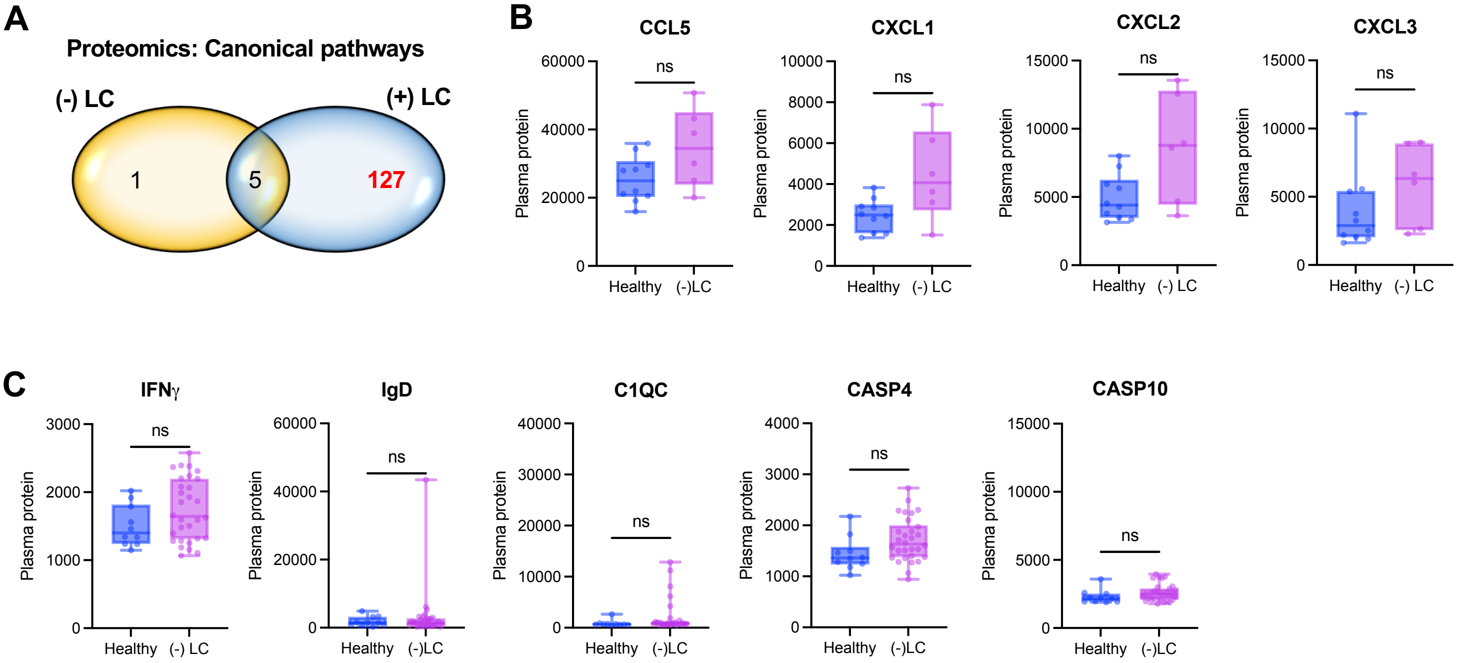

### Figure S2

**Fig. S2. Nitric oxide biosynthesis in COVID patients.**

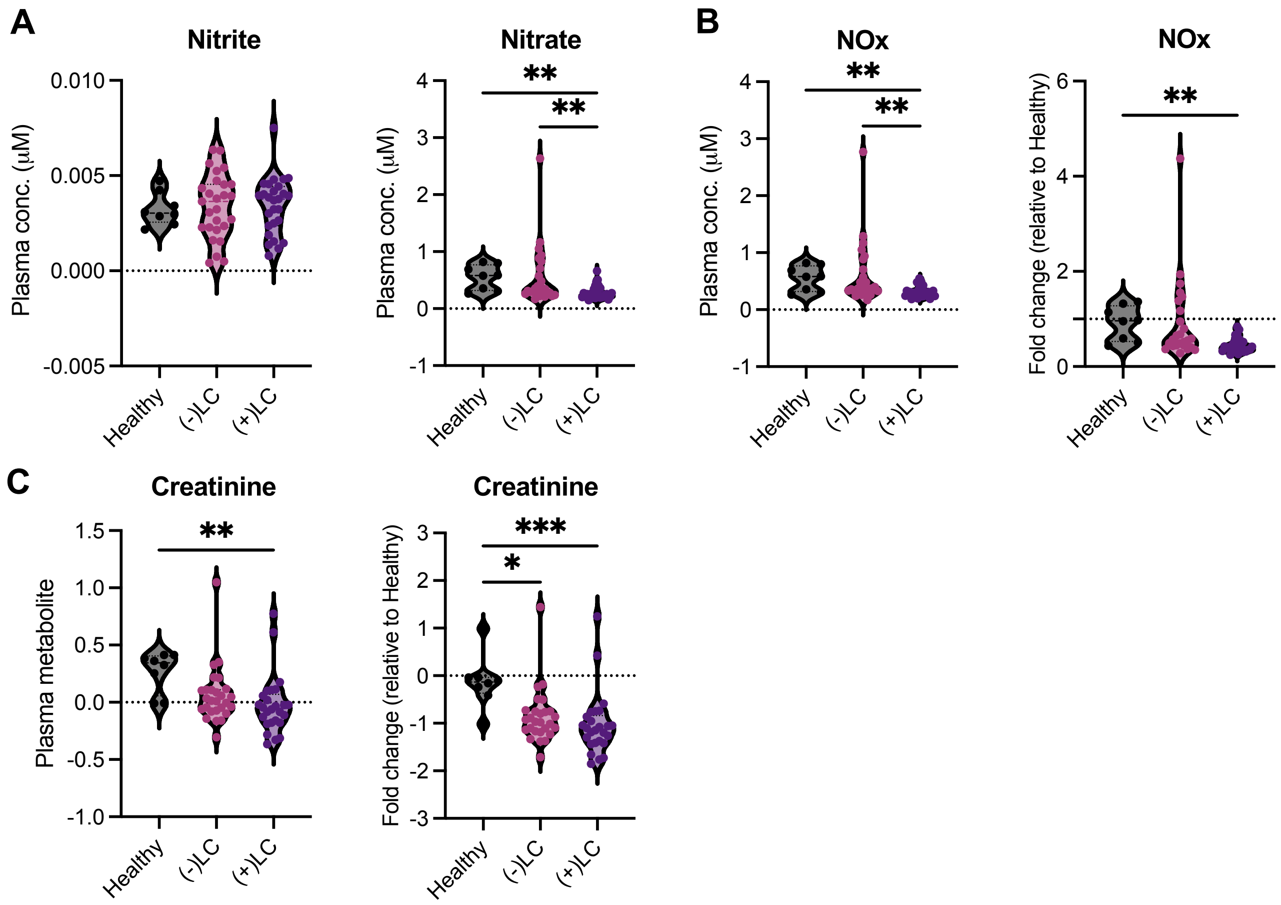
