## Supplementary material for "Immunometabolic rewiring in long COVID patients with chronic headache": Table S1

**Table S1. Overview of the clinical demographics and parameters of COVID19 patients upon COVID19 diagnosis.**

| | COVID-negative | No Long-COVID | Long-COVID-headache |
|---|---|---|---|
| **Demographics: Race (%)** | | | |
| Asian | 0 | 0 | 0 |
| Black | 0 | 17 | 20 |
| Other | 0 | 14 | 0 |
| White | 100 | 69 | 80 |
| Male | 20 | 44 | 19 |
| Age (yrs) | 63.1 | 55.3 | 42.7 |
| **Risk factors: Smoking (%)** | | | |
| Current Smoker | 0 | 7 | 18 |
| Former Smoker | 75 | 53 | 9 |
| Non-smoker | 25 | 40 | 64 |
| Unknown smoking status | 0 | 0 | 9 |
| **COVID symptoms presented upon COVID19 diagnosis:** | | | |
| Cough (%) | 75 | 73 | 82 |
| Fever (%) | 25 | 36 | 36 |
| Fatigue (%) | 25 | 63 | 55 |
| Sputum production (%) | 0 | 30 | 9 |
| Flu like symptoms (%) | 0 | 60 | 73 |
| Diarrhea (%) | 25 | 30 | 18 |
| Loss of appetite (%) | 0 | 30 | 27 |
| Vomiting (%) | 50 | 13 | 36 |
| **Co-morbidities:** | | | |
| COPD emphysema (%) | 25 | 11 | 0 |
| Asthma (%) | 50 | 20 | 9 |
| Diabetes (%) | 75 | 40 | 19 |
| Hypertension (%) | 100 | 57 | 27 |
| Coronary artery disease (%) | 75 | 17 | 0 |
| Heart failure (%) | 25 | 14 | 0 |
| Cancer (%) | 25 | 9 | 0 |
| Transplant history (%) | 25 | 0 | 0 |
| Multiple sclerosis (%) | 0 | 0 | 9 |
| Connective tissue disease (%) | 25 | 9 | 0 |
| Inflammatory bowel disease (%) | 25 | 6 | 0 |
| **Laboratory values upon COVID19 diagnosis:** | | | |
| Platelets (k/uL) | 155.5 | 211.5 | 194.8 |
| Hematocrit (%) | 39.7 | 37.5 | 40.4 |
| WBC (k/uL) | 4.9 | 6.4 | 5.1 |
| BUN (mg/dL) | 19.5 | 18.3 | 11.4 |
| Creatinine (mg/dL) | 1.5 | 1.3 | 0.91 |
| AST (U/L) | 33 | 32.3 | 31.8 |
| CRP | - | 7.8 | 2 |
| **Baseline medication use at time of COVID diagnosis:** | | | |
| NSAIDS (%) | 0 | 23 | 18 |
| Steroids (%) | 25 | 23 | 9 |
| Melatonin (%) | 0 | 7 | 0 |
