## Supplementary material for "Immunometabolic rewiring in long COVID patients with chronic headache": Table S2

**Table S2. Canonical pathways that are uniquely altered in (-)LC patients. For Fig. 3.**

| Timepoint | Canonical Pathways | z-score: (-)LC | z-score: (+)LC |
|---|---|---|---|
| >-8 days | GADD45 Signaling | 2.236 | 0 |
| | Regulation of Actin-based Motility by Rho | -2.111 | N/A |
| | Melanocyte Development and Pigmentation Signaling | -2.111 | N/A |
| | ERK5 Signaling | -2.138 | N/A |
| | Retinoic acid Mediated Apoptosis Signaling | -2.236 | 1.633 |
| | Remodeling of Epithelial Adherens Junctions | -2.236 | N/A |
| | HMGB1 Signaling | -2.324 | 1.897 |
| | NGF Signaling | -2.324 | 1.633 |
| | Death Receptor Signaling | -2.324 | 1.134 |
| | Role of IL-17F in Allergic Inflammatory Airway Diseases | -2.333 | 1 |
| | Gαs Signaling | -2.333 | 0.816 |
| | Ephrin Receptor Signaling | -2.524 | 1.89 |
| | White Adipose Tissue Browning Pathway | -2.646 | -0.302 |
| | Th1 Pathway | -3.317 | 0.707 |
| -7-7 days | Signaling by Rho Family GTPases | 2.333 | 1.633 |
| | Interferon Signaling | 2.236 | N/A |
| | Kinetochore Metaphase Signaling Pathway | 2.121 | N/A |
| | Estrogen-mediated S-phase Entry | 2 | N/A |
| | Senescence Pathway | -2 | -0.816 |
| | RHOGDI Signaling | -2.449 | -1.633 |
| 8-30 days | Cyclins and Cell Cycle Regulation | 2.449 | N/A |
| | EIF2 Signaling | 2.324 | N/A |
| | Non-Small Cell Lung Cancer Signaling | -2 | N/A |
| | Chondroitin Sulfate Degradation (Metazoa) | -2 | N/A |
| | ERK5 Signaling | -2.121 | N/A |
| | UVA-Induced MAPK Signaling | -2.121 | N/A |
| | NF-κB Signaling | -2.138 | N/A |
| | Type II Diabetes Mellitus Signaling | -2.333 | N/A |
| | G-Protein Coupled Receptor Signaling | -2.405 | -1 |
| | Lymphotoxin β Receptor Signaling | -2.449 | N/A |
| | Dendritic Cell Maturation | -2.5 | -0.447 |
| | Melanocyte Development and Pigmentation Signaling | -2.53 | N/A |
| 1-3 months | Breast Cancer Regulation by Stathmin1 | -2.111 | -0.577 |
| >3 months | N/A | N/A | N/A |
